## Supplementary Figures for "North American deer mice are susceptible to SARS-CoV-2"

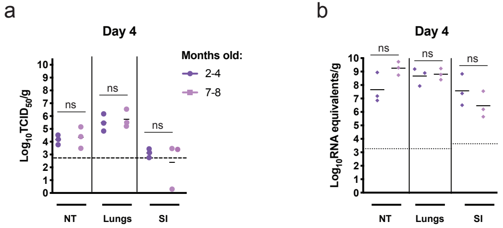


**Extended Data Fig. 1: Viral burden in the tissues by age.** Eight to thirty two-week old male or female deer mice (*P. maniculatus*) were inoculated with $1x10^5$ $TCID_{50}$ of SARS-CoV-2 by an intranasal route (i.n.) of administration. **a**, Infectious viral load in the nasal turbinates, lung, and proximal small intestine at 4 dpi. **b**, Viral RNA load in the nasal turbinates, lung, and proximal small intestine at 4 dpi. Bars indicate means. Dashed lines and dotted lines indicate the limit of detection for the $TCID_{50}$ assay and qRT-PCR assay, respectively. ns = $P > 0.05$, ANOVA with multiple comparison correction.

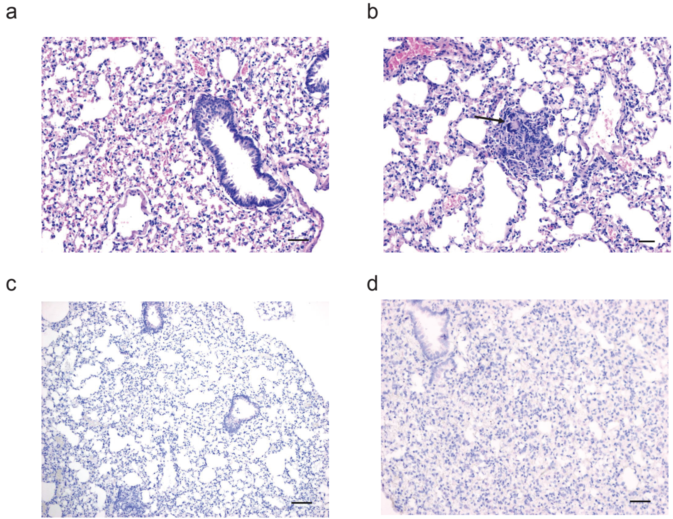


**Extended Data Fig. 2: Histological analysis of lungs at 21 days following SARS-CoV-2 exposure.** Adult male and female deer mice were exposed to $10^5$ $TCID_{50}$ SARS-CoV-2 by an i.n. route of infection. Lesions were not observed at this time point (**a**) except for a small focus of inflammation (**b**) with presence of syncytial cell (arrow). ISH using both anti-sense probe (**c**) and sense probe (**d**) was negative. Scale bars = 50 µm (a-b) and 100 µm (c-d).

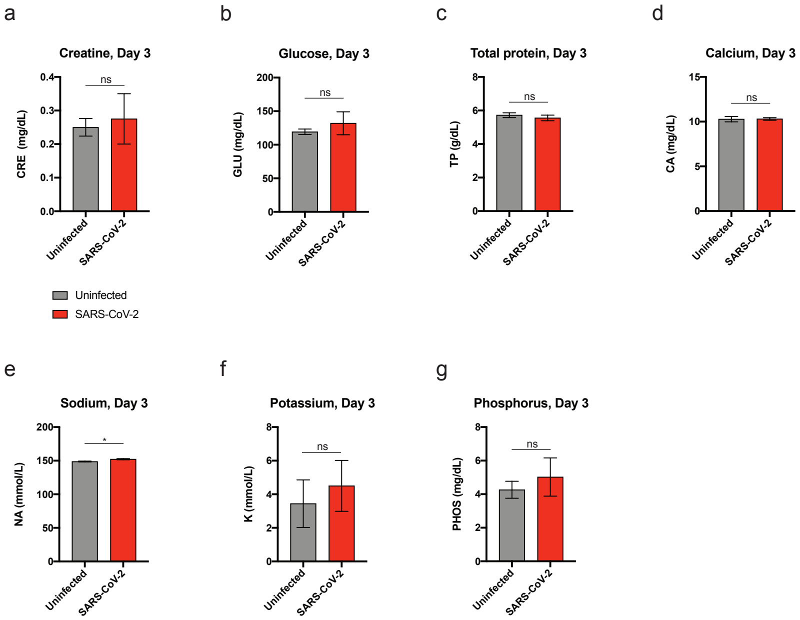


**Extended Data Fig. 3. Serum biochemical values.** Serum biochemistry were measured in uninfected and SARS-CoV-2-infected deer mice at 3 dpi ($10^6$ $TCID_{50}$, i.n. route). Error bars represent SEM. * = $P < 0.05$, ns = $P > 0.05$, unpaired t test with Welch's correction.

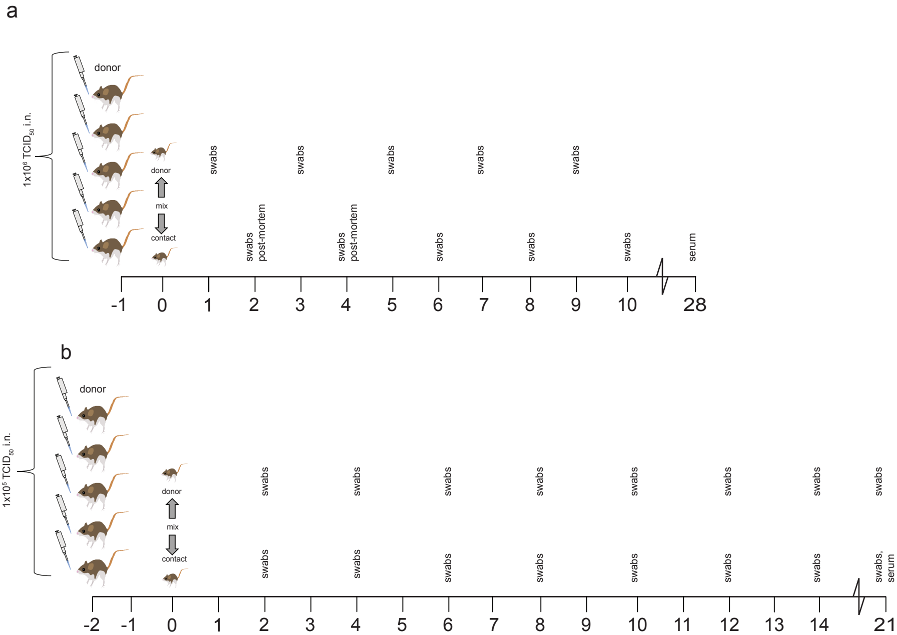


**Extended Data Fig. 4. Schematic of deer mouse SARS-CoV-2 transmission study 1 & 2.** Adult male and female deer mice were exposed to **a**) $1x10^6$ $TCID_{50}$ or **b**) $1x10^5$ $TCID_{50}$ SARS-CoV-2 by an i.n. route of infection. At **a**) 1 dpi or **b**) 2 dpi individual inoculated donor deer mice were transferred to a new cage and co-housed with a single naïve deer mouse (1:1 ratio) to assess SARS-CoV-2 transmission by direct contact. Deer mice were either maintained in direct contact throughout the study for serial swabbing or were humanely euthanized on 2 or 4 dpi for tissue collection.
